# Integration of cell cycle signals by multi-PAS domain kinases

**DOI:** 10.1101/323444

**Authors:** Thomas H. Mann, Lucy Shapiro

**Affiliations:** Department of Developmental Biology, Stanford University School of Medicine Stanford, California, 94305

## Abstract

Spatial control of intracellular signaling relies on signaling proteins sensing their subcellular environment. In many cases, a large number of upstream signals are funneled to a master regulator of cellular behavior, but it remains unclear how individual proteins can rapidly integrate a complex array of signals within the appropriate spatial niche within the cell. As a model for how subcellular spatial information can control signaling activity, we have reconstituted the cell pole-specific control of the master regulator kinase/phosphatase CckA from the asymmetrically dividing bacterium *Caulobacter crescentus.* CckA is active as a kinase only when it accumulates within a microdomain at the new cell pole, where it co-localizes with the pseudokinase DivL. Both proteins contain multiple PAS domains, a multifunctional class of sensory domains present across the kingdoms of life. Here, we show that CckA uses its PAS domains to integrate information from DivL and on its own oligomerization state to control the balance of its kinase and phosphatase activities. We reconstituted the DivL-CckA complex on liposomes *in vitro* and found that DivL directly controls the CckA kinase-phosphatase switch, and that stimulation of either CckA catalytic activity depends on the second of its two PAS domains. We further show that CckA oligomerizes through a multi-domain interaction that is critical for stimulation of kinase activity by DivL, while DivL stimulation of CckA phosphatase activity is independent of CckA homo-oligomerization. Our results broadly demonstrate how signaling factors can leverage information from their subcellular niche to drive spatiotemporal control of cell signaling.

**Significance:** Cells must constantly make decisions involving many pieces of information at a molecular level. Kinases containing multiple PAS sensory domains detect multiple signals to determine their signaling outputs. In the asymmetrically dividing bacterium *Caulobacter crescentus,* the multi-sensor proteins DivL and CckA promote different cell types depending upon their subcellular location. We reconstituted the DivL-CckA interaction *in vitro* and showed that specific PAS domains of each protein function to switch CckA between kinase and phosphatase activities, which reflects their functions *in vivo.* Within the context of the cell, our reconstitution illustrates how multi-sensor proteins can use their subcellular location to regulate their signaling functions.

## Introduction

Asymmetric cell division is a fundamental mechanism for generating cell type diversity across the kingdoms of life. Accomplishing an asymmetric division requires coordination between cell cycle-dependent gene expression and the dynamic subcellular localization and function of signaling proteins (1). A well-studied model system exhibiting asymmetric division is the bacterium *Caulobacter crescentus*, which generates two distinct daughter cells every cell division (Fig. 1A). A master regulator of cellular identity is the transcription factor CtrA, which when phosphorylated, directly controls gene expression of over 90 cell cycle regulated promoters while also inhibiting the initiation of DNA replication (2–4). CtrA~P is present and active as a transcription factor in the motile, replication-incompetent swarmer cell. Dephosphorylation and proteolysis of CtrA~P permits differentiation into a sessile stalked cell and the beginning of DNA replication (5). Upon progression into the predivisional stage, CtrA proteolysis ceases and the transcription of *ctrA* is activated. Concurrently, a set of signaling proteins localize to the new cell pole, opposite the stalk, to promote CtrA phosphorylation and the biogenesis of the flagellum and pili. Spatial and temporal control of CtrA~P thus coordinates cell type identity with DNA replication and cell cycle progression.

**Figure 1.**
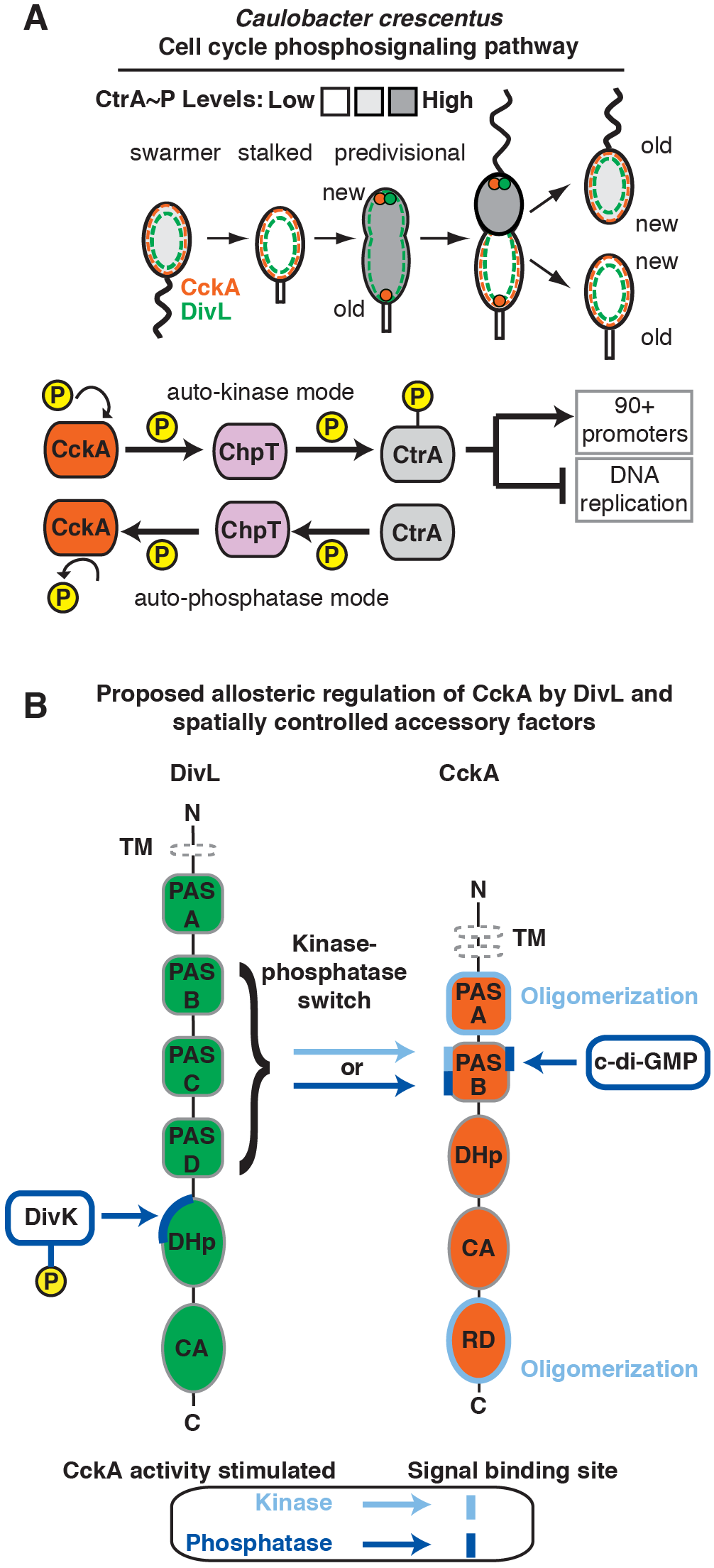
DivL regulates CckA kinase and phosphatase activities as a function of the cell cycle. A. DivL (green) and CckA (orange) subcellular localizations are dynamic over the *Caulobacter* cell cycle. In the swarmer and stalked cells, both proteins are diffusely deployed around the membrane (dashed lines). In the predivisional cell, DivL accumulates at the new cell pole (green circle), where it is essential for both CckA new pole-localization and CckA kinase activity. CckA (orange circles) accumulates at both the old and new cell poles, and both proteins maintain a partial diffuse population outside of the polar niches. CckA kinase and phosphatase activities control the cell cycle dependent phosphorylation state of the master regulator, CtrA. Phosphorylated CtrA~P (grayscale) inhibits the initiation of DNAreplication and activates transcription of over 90 promoters. Phosphate is shuttled between CckA and CtrA via the transfer protein ChpT (lavender). B. CckA and DivL both integrate signals to determine CckA kinase activity. Light blue arrows represent transmission of a CckA kinase-stimulating signal, whereas dark blue arrows represent a phosphatase-stimulating signal. Light and dark bars represent the binding sites for those signals, respectively. DivK~P acts as an accessory factor to DivL, binding in the DHp domain and reconfiguring the DivL PAS domains signaling to promote CckA phosphatase activity. Stimulation of CckA kinase and phosphatase activities by DivL are both transmitted in a CckA PAS-B dependent manner, likely through different binding sites. In addition to receiving a signal from DivL, CckA oligomerization and c-di-GMP promote CckA kinase and phosphatase activities, respectively. CckA catalytic output is determined upon integration of all available signals. TM represents the transmembrane tethers of the two proteins, replaced by His-tags in our *in vitro* experiments.

The bifunctional histidine kinase/phosphatase CckA controls the phosphorylation state of CtrA (6–9). CckA uses sensory PAS domains to change its activity depending upon its subcellular location and the progression of the cell cycle (10, 11). We previously reconstituted CckA on liposomes *in vitro* to show that its kinase activity depends upon CckA surface density, and that this density-dependent response requires the first of its two PAS domains (11). In predivisional cells, CckA is a kinase when it accumulates at high density at the new cell pole, supplying the phosphate for CtrA~P through a phosphorelay protein, ChpT (Fig 1A) (12). However, in the stalked cell, high levels of the second messenger c-di-GMP (cdG) promote CckA phosphatase activity, which depletes the CtrA pathway of phosphate (Fig 1B) (11, 13,14).

Upstream of CckA, the pseudo-histidine kinase DivL is necessary for CckA localization to the new cell pole and essential for CckA kinase activity in predivisional cells (10, 15). CckA co-immunoprecipitates with DivL, suggesting that CckA regulation requires complex formation with DivL (10). DivL is also essential for effective inhibition of CckA kinase activity when the two proteins are away from the new cell pole. *In vivo,* interaction of DivL and the phosphorylated response protein DivK~P leads to inhibition of CckA kinase activity (16, 17). A tyrosine replaces histidine at the active site of the DivL pseudokinase with no apparent catalytic activity (18). It has been suggested that DivL switches between conformations that promote or inhibit CckA kinase activity when the two are in a complex (17).

In most histidine kinases, extracellular or intracellular signals interact with N-terminal PAS domains leading to a conformational change that in turn regulates the activity of the catalytic domains (19, 20). Kinases can also use this conserved structural link in the reverse direction. DivL (Fig 1B) uses its kinase-like domain to bind to the phosphorylated response regulator DivK~P (17), predicted to drive a conformational change within DivL’s PAS domains. This rearrangement in the PAS domains is then propagated as a regulatory signal to CckA’s own PAS domains, regulating CckA catalytic activity (Fig 1B) (17), and previous studies indicate that the PAS domains of the two proteins are sufficient for communication between the two proteins (10, 21). However, it has remained unclear whether DivL directly regulates CckA catalytic activity, and how a multi-PAS domain protein can pass distinct signals to a target kinase.

Here, we show that DivL directly regulates the kinase-phosphatase switch of CckA *in vitro* when the two proteins are reconstituted on proteoliposomes. DivL can inhibit CckA kinase activity in the absence of any ligands, instead stimulating CckA phosphatase activity. We also show that a point mutant in DivL can constitutively activate CckA kinase activity, and that stimulation of either kinase or phosphatase activity requires the PAS domains of DivL and PAS-B of CckA. Additionally, oligomerization of CckA is critical for DivL stimulation of kinase activity but not its stimulation of CckA phosphatase activity. While we must keep in mind that this reconstitution approach represents a simplified system that lacks additional factors present *in vivo*, our previous experiments using reconstituted CckA on liposomes have been supported by subsequent work demonstrating agreement that the CckA surface density that is critical for kinase activity in our *in vitro* experiments matches the surface density of CckA present at the new cell pole *in vivo* (22). In this study, we reconstitute an additional aspect of subcellular regulation of CckA. We propose that DivL integrates information about subcellular localization and cell cycle progression to toggle CckA between its kinase and phosphatase modes in a PAS domain-dependent manner (Fig 1B).

## Results

### DivL directly inhibits CckA kinase activity

We set out to test whether DivL can directly regulate CckA catalytic activity *in vitro*. Membrane tethering is essential for CckA polar localization and kinase activity *in vivo* (6), suggesting that that membrane attachment may be important for productive interaction between CckA and DivL (6). We purified CckA and DivL constructs containing N-terminal His-tags in place of their transmembrane helices, and we tethered the two proteins to Ni-NTA groups on a fluid lipid surface (Fig 2A) (11). Unless otherwise stated, we present data using a DivL construct lacking its N-terminal domain A, which has distant homology to PAS domains (23, 24), due to the increased protein stability of the shortened construct and its indistinguishable behavior from the full length construct in all of our assays. To probe DivL’s regulation of CckA kinase activity, we incubated CckA and DivL at equimolar concentrations of 5 μM either in solution or an equivalent amount of protein at 350 molecules of each protein per liposome. CckA kinase activity was measured by auto-phosphorylation in the presence of [γ-P32]-ATP followed by autoradiography. Strikingly, co-loading DivL with CckA on liposomes inhibited CckA kinase activity, whereas co-incubation of the two proteins at 5 μM each in solution, with no liposomes, yielded no change to kinase activity (Fig 2B), suggesting that membrane tethering is critical for interaction between the two proteins.

**Figure 2.**
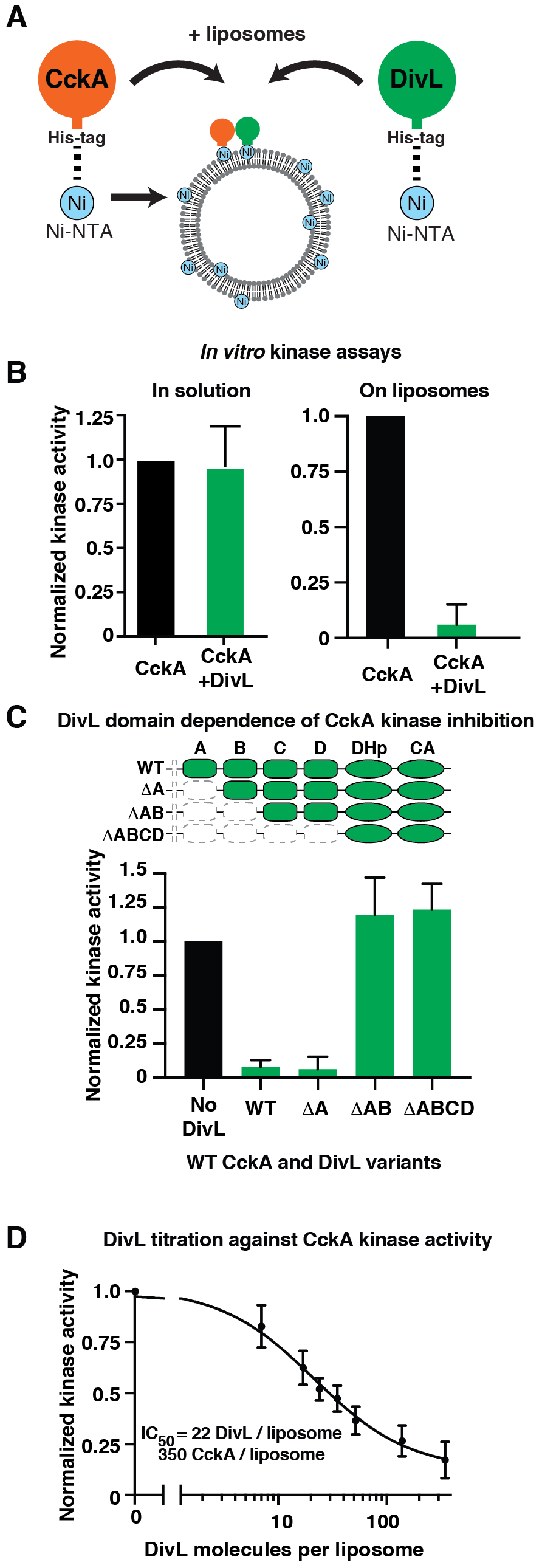
DivL directly regulates CckA kinase activity in a DivL PAS domain-dependent manner. A. Purified CckA and DivL can be tethered to liposomes to facilitate their interaction. The liposomes are doped with Ni-NTA-containing lipids, and the proteins are attached via N-terminal His-tags, mimicking their membrane tethers *in vivo.* B. CckA kinase activity was measured by allowing CckA to autophosphorylate for 3 minutes in the presence of [γ-P32]-ATP. The effect of DivL on CckA kinase activity was measured either free in solution or on liposomes. C. CckA kinase activity was measured in the presence of different DivL PAS domain truncations.For each DivL variant, removal of a domain is represented by the dashed, empty boxes, while presence of the domain is represented by green shading. D. CckA autophosphorylation was measured in the presence of varying surface densities of DivL. Kinase activity is normalized to the no-DivL condition in each panel. All protein densities except forthe DivL titration in panel D were 350 molecules per liposome. Error bars represent the standard deviation of at least 3 experiments for all panels.

Previous studies indicated that the PAS domains of CckA and DivL (Fig 1B) are important for both proteins’ signaling functions (10, 11, 17). To test whether the DivL PAS domains are critical for its inhibition of CckA autophosphorylation, we purified a series of DivL PAS domain deletion mutants, progressively truncating PAS domains from the N-terminus (17). We then incubated these DivL PAS domain truncations with CckA on liposomes to measure their effects on CckA kinase activity (Fig 2C). While the full-length DivL and a construct lacking PAS domain A inhibited CckA to the same extent, the DivL constructs ΔPAS-AB and ΔPAS-ABCD did not impact CckA kinase activity, demonstrating that the two N-terminal PAS domains of DivL were necessary for a functional interaction on the membrane. This finding is consistent with the prior model suggesting that DivL and CckA communicate through contacts in their PAS domains (17).

To determine the potency of DivL as an inhibitor of kinase activity, we titrated DivL at different surface densities against a fixed density of CckA on liposomes. DivL robustly inhibited CckA kinase activity with an IC_50_ of 22 DivL molecules per liposome (95% confidence interval: 15-31 DivL/liposome), 15-fold lower than the CckA surface density (Figure 2D). This sub-stoichiometric inhibition implies that DivL inhibition of kinase activity may be due to enhanced phosphatase activity by DivL or CckA towards CckA~P. These *in vitro* data indicate that in the absence of accessory factors, DivL can sub-stoichiometrically inhibit CckA kinase activity (Fig 2D). *In vivo*, DivL binds the accessory factor DivK~P to inhibit CckA kinase activity when these proteins are away from the new cell pole (16), while DivL is critical for stimulation of CckA kinase activity when the two proteins co-localize at the new cell pole (10, 16). Based on these findings, we propose that DivL binds another accessory factor at the new cell pole to facilitate its stimulation of CckA kinase activity.

### DivL stimulates CckA auto-phosphatase activity

Inhibition of kinase activity in many HKs frequently represents a concerted switch to a conformation that has increased phosphatase activity towards the cognate receiver domain (25, 26). It was previously shown that binding of the second messenger (cdG) causes CckA to switch from kinase mode to phosphatase mode (11, 13, 27). This finding suggests that DivL’s inhibition of CckA kinase activity may, similar to cdG, induce a switch from a kinase-active conformation to a phosphatase-active one. Alternatively, we considered the possibility that DivL may directly perform the phosphatase activity on CckA’s receiver domain.

We therefore tested whether DivL promotes CckA dephosphorylation *in vitro*, monitoring the rate at which purified CckA~P lost phosphate when incubated with or without DivL on liposomes (Fig 3A). The presence of DivL on liposomes with CckA stimulated the loss of CckA~P signal, indicating that one of the proteins was performing phosphatase activity (Fig 3B). DivL did not stimulate phosphate decay from the phosphatase-deficient mutant CckA V366P (28),indicating that CckA, and not DivL, is responsible for performing phosphatase activity (Fig. S1A, B). To test if the CckA PAS domains are necessary for this response to DivL (Fig 1B), we repeated the phosphatase experiment for the CckA variants ΔPAS-A, ΔPAS-B, and ΔPAS-AB. CckA ΔPAS-A rapidly lost phosphate upon co-loading with DivL on liposomes, indicating that CckA PAS-A is not necessary for interaction with DivL or phosphatase activity (Fig 3C).

**Figure 3.**
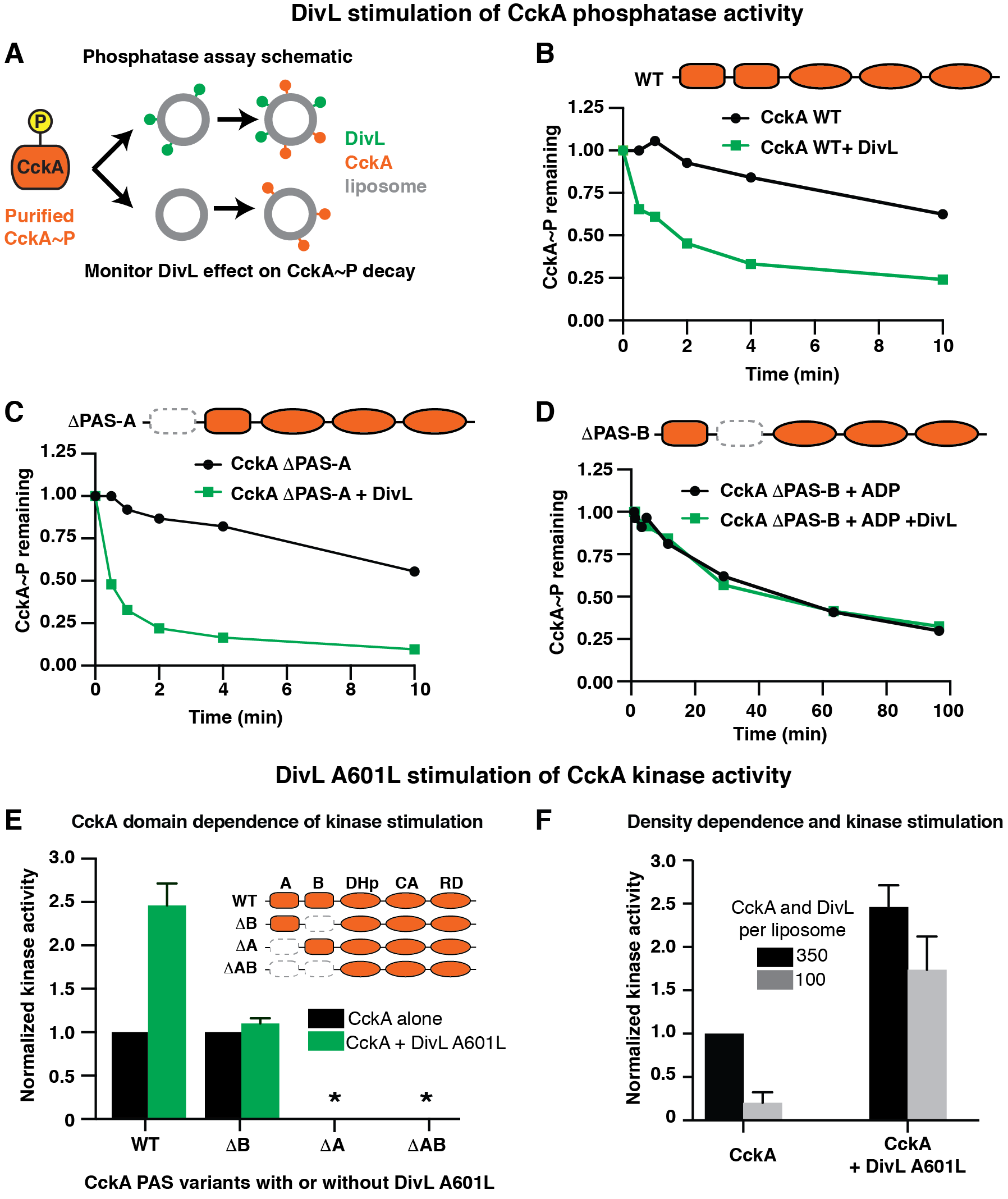
DivL can stimulate CckA as an auto-phosphatase or as an auto-kinase, in a CckA PAS-B dependent manner. A. A schematic illustrates how to test whether DivL affects CckA phosphatase activity *in vitro.* More details are available in the Methods section. B. The loss of CckA~P signal following ATP depletion was monitored by quenching reaction aliquots at 0.5, 1, 2, 4, and 10 minutes following mixing of CckA and DivL. (C, D) The experiment was repeated for the CckA variants CckA ΔPAS-A and ΔPAS-B, respectively. The experiment was modified for CckA ΔPAS-B by omitting the desalting column step, keeping 500 μM ADP in solution following hexokinase treatment, and monitoring CckA~P for approximately 100 minutes. PAS domain deletions are represented as empty, dashed boxes. E. The CckA PAS domain dependence for autokinase activation by DivL A601L was tested. The kinase activities of CckA variants ΔPAS-A, ΔPAS-B, and ΔPAS-A-B (orange domain schematics) were tested with DivL A601L (green) or without it (black) at350 molecules per liposome for each protein. Kinase activity is normalized to the CckA-only condition for each variant. For constructs lacking PAS-A, kinase activity at this density was very low and is represented by an *. F. The density dependence ofCckA response to DivL A601L was tested. In each experiment, CckA was present at 5 μM total in solution, with liposomes added to bring CckA to either 350 (black) or 100 (gray) molecules per liposome. DivL A601L was present on liposomes in equimolar quantities to each CckA condition where applicable. For panels B-D, CckA~P decay traces are representative of at least two independent experiments, and for panels E andF, error bars represent the range of at least two experiments.

Strikingly, CckA constructs lacking PAS-B did not show a change in autophosphatase activity in the presence of DivL (Fig 3D, Fig. S1C). Because their autophosphatase activity was reduced compared to WT CckA, we took advantage of the finding that ADP binding promotes the phosphatase-competent conformation of histidine kinases (25, 27) to show that CckA ΔPAS-B and ΔPAS-AB mutants do retain phosphatase function (Fig. S2). Thus, CckA PAS-B is necessary for auto-phosphatase stimulation by DivL. It is formally possible that CckA constructs lacking PAS-B are conformationally restricted in a manner that prevents interaction between DivL and another part of CckA. Given that the N-terminus of CckA comprising its transmembrane region, PAS-A, and PAS-B together are sufficient for response to DivL *in vivo* though (10), and PAS-A is not necessary for response to DivL in this assay (Fig 3E), we propose that PAS-B is the CckA domain that communicates with DivL.

We previously showed that PAS-B of CckA is necessary for binding to cdG, and that it is sufficient for binding when domains involved in dimerization and oligomerization are also present (11). Because our findings indicate that DivL, like cdG, also signals through CckA’s PAS domains, we further tested whether cdG cooperatively inhibited CckA in the presence of DivL. While cdG inhibition has been shown to be cooperative with ADP binding (27), we observed additive rather than cooperative inhibition of CckA kinase activity by cdG and DivL inhibition of CckA kinase activity (Fig. S1D).

### A DivL point mutant directly stimulates CckA kinase activity

Prior studies indicated that DivL switches between activating or inhibiting CckA kinase activity *in vivo,* depending on the stage of the cell cycle (10, 16), and uses accessory factors such as the phosphorylated response regulator DivK~P to determine its activity towards CckA.

Tsokos et al. found that a DivL point mutant, A601L, enhanced CckA kinase activity by fivefold *in vivo* during the stalked cell phase of the cell cycle even when the proteins were localized away from the new cell pole (16). This finding suggests that DivL A601L activates CckA kinase activity independent of upstream signals. Moreover, structure-function analysis suggested that the A601L mutation changes the global conformation of DivL in a PAS domain-dependent manner (17).

To determine if DivL A601L directly stimulates CckA kinase activity through the CckA PAS domains. The CckA PAS domain mutants had different capacities for response to DivL A601L (Fig 3E). While CckA ΔPAS-B retained kinase activity when tethered to liposomes, its kinase activity did not respond to the presence of DivL A601L. Conversely, CckA variants lacking PAS-A did not demonstrate significant kinase activity on liposomes as we previously observed, and DivL A601L did not rescue their kinase activities. These data indicate that PAS-B is specifically required for a kinase response to DivL A601L, while PAS-A is critical for CckA kinase activity. Because PAS-A is critical for CckA surface density-dependent kinase activity, we tested compared the effects of DivL A601L stimulation of CckA kinase activity at different densities (Fig 3F). Even at reduced surface density, DivL A601L greatly stimulated CckA kinase activity, albeit to a lower total extent than when the two proteins were incubated at high density. This finding is consistent with the finding that A601L promotes CckA kinase activity even when the proteins are diffusely localized *in vivo* (16). Altogether, these data indicate that offer proof-of-principle that DivL can stimulate CckA kinase activity in a PAS domain specific manner and suggest that additional upstream signals may push DivL into this kinase activity promoting state*in vivo.*

### CckA oligomerizes through PAS-A and its receiver domain

The requirement of high surface densities for CckA kinase activity on liposomes suggests that it may need to oligomerize to become active as a kinase (11). While the catalytic core of CckA, containing the DHp and CA domains, forms a canonical *trans*-phosphorylating dimer (27), we hypothesized that its PAS and receiver domains may drive higher oligomerization. There are a handful of histidine kinases believed to function as tetramers, including the kinase RegB, which loses kinase activity upon tetramerization (29–31). We assayed CckA oligomerization in solution using analytical size exclusion chromatography (SEC) to predict the molecular mass of CckA complexes. Indeed, we found that WT CckA eluted as a single tetrameric peak when injected onto the column at approximately 100 μM in solution (Fig 4A, Table S1). Knowing that PAS domains frequently mediate oligomerization (19), and that PAS-A is critical for density-dependent kinase activity, we tested the PAS domain dependence of CckA oligomerization. Removal of CckA PAS-A showed a reduction in oligomerization, with a peak center at 3.1 CckA per complex, indicating that PAS-A contributes to higher oligomerization, but that another domain in the protein continues to drive oligomerization in the absence of PAS-A (Fig 4B). Removal of PAS-B yielded a smaller effect on CckA oligomerization than PAS-A (Supplementary Table S1).

**Figure 4.**
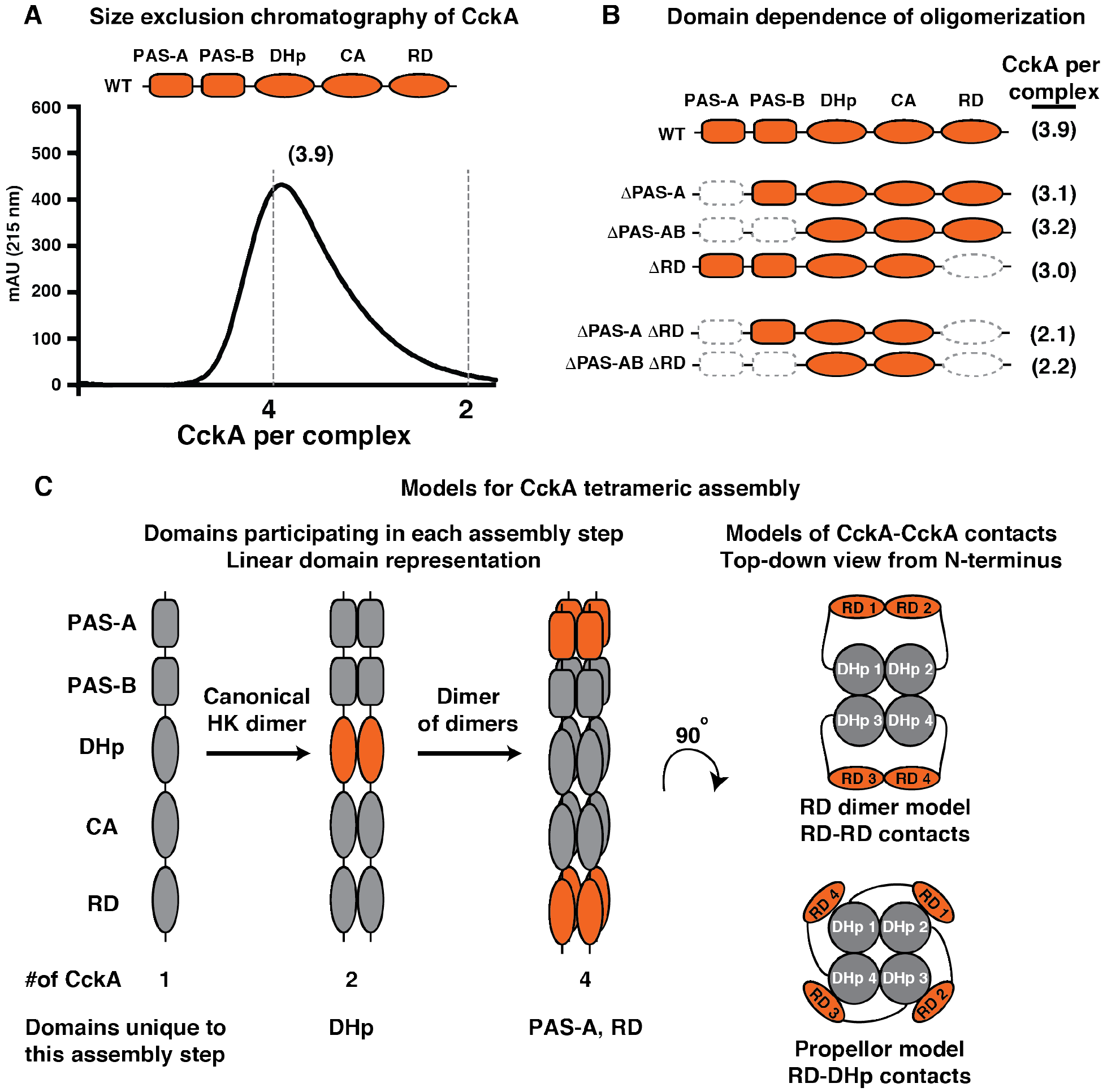
CckA oligomerizes through a multi-domain interaction. A. The oligomerization state of CckA in solution was measured by analytical size exclusion chromatography. The elution trace from the size exclusion column is shown, with dashed vertical lines indicating the predicted elution volumes for a tetramer or dimer. Predicted molecular mass of the CckA complex was interpolated using a standard curve of known protein masses. B. The size exclusion experiment was repeated for the CckA variants APAS-A, ΔPAS-A-B, ARD, ΔPAS-A-ARD, and ΔPAS-A-B-ARD. The projected CckA per complex is shown to the right of each variant. A complete list of SEC experiments is given in Supplementary Table S1. In each experiment, CckA was loaded onto the column at a concentration of roughly 100 μM. A complete list of SEC experiments is given in Supplementary Table S1. C. (Left) A two-step model for CckA oligomerization illustrates the domains involved in each oligomerization step, highlighted in orange. The DHp domain promotes a conserved dimerization step. PAS-A and the RD are both critical for assembly into a dimer of dimers. This representation does not reflect the relative positions of the domains in 3D. (Right) Two possible models for how RD contributes to CckA oligomerization are shown as 2D projections looking down the central DHp axis from the N-terminus, one by RD-RD dimerization, common among response regulators, or by RD-DHp contacts that bridge two dimers.

Because the DHp domain typically only provides a conserved dimerization interface, we tested whether the receiver domain of CckA might also contribute to tetramerization. CckA DRD indeed showed a partial reduction in oligomerization (Fig 4B), as well as a reduction in kinase activity on liposomes (Fig. S3). Simultaneous deletion of CckA variants lacking both PAS-A and RD (ΔPAS-A DRD and ΔPAS-A-B DRD), resulted in CckA attaining only a dimeric state. Given that the catalytic core of CckA has been shown to form a canonical dimer (27), it seems apparent that the dimers should not have to be disassembled and reassembled into a fourfold-symmetric tetramer. Similarly, we hypothesize that the elution peak centered at approximately three CckA molecules/complex represents an exchange between states containing two and four CckA molecules/complex and tetrameric states, rather than forming a distinct trimer. We propose that CckA’s PAS-A and the RD mediate the assembly of conserved HK dimers into a dimer of dimers(Fig 4C).

## Discussion

Cells must constantly integrate information to coordinate cell cycle events. For processes that require spatial control, such as differentiation and asymmetric division,their signaling proteins must additionally be able to recognize and respond to upstream factors found at distinct subcellular locations. Multi-sensor histidine kinases constitute a large set of signaling proteins which have the potential to respond to multiple signals, enabling the processing of information within subcellular niches. In this study, we have shown that the pseudokinase DivL controls the kinase-phosphatase switch of CckA through the PAS domains of the two proteins. DivL stimulation of kinase activity further requires CckA homo-oligomerization, illustrating how subcellular accumulation of CckA at the cell pole can be used as input information for the regulation of cellular asymmetry (Fig. 5).

**Figure 5.**
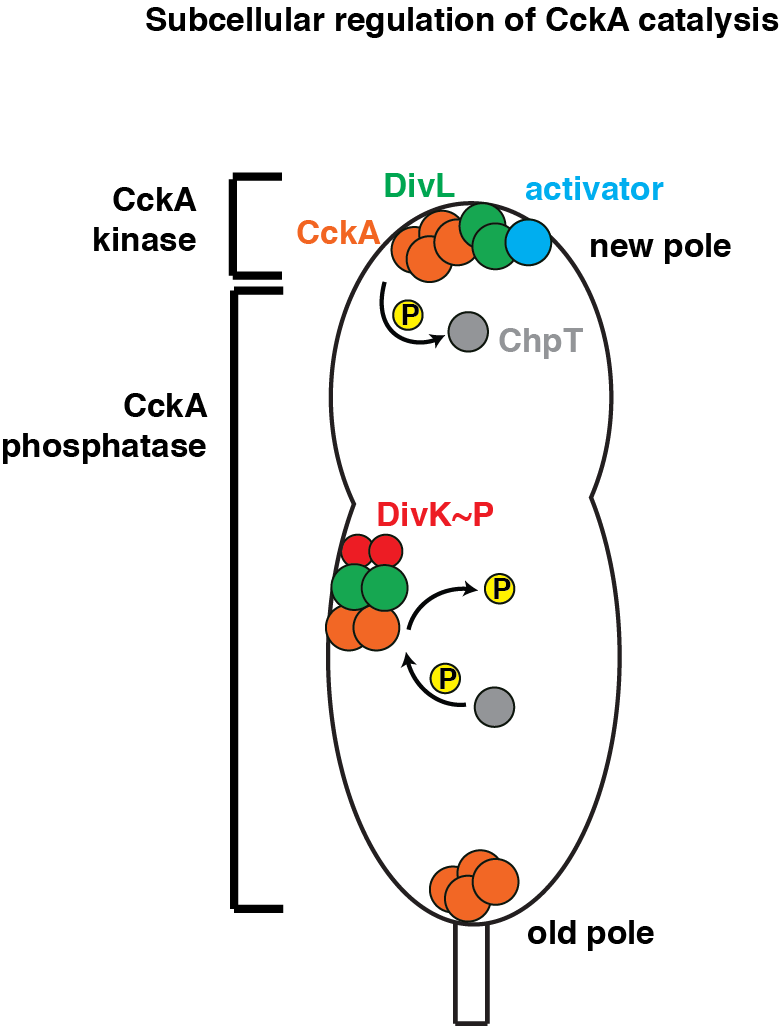
Multiple signals converge on CckA to regulate its kinase-phosphatase switch depending upon its subcellular niche. A model of spatially controlledkinase/phosphatase signaling. CckA (orange) accumulates at the new cell pole, favoringoligomerization and formation of a kinase-active complex with DivL (green), possiblywith thehelp of an additional kinase-stimulating factor (blue) specifically localized at that pole. Kinase activity leads to phosphorylation of ChpT (gray) and eventually CtrA. Outside of the new pole the DivL/DivK~P (red) complex can promote CckA phosphatase activity regardless of CckA’s oligomeric state, siphoning phosphate out ofthe CtrA pathway.

We found that DivL can directly stimulate both CckA kinase and phosphatase activities in a PAS-B dependent manner (Fig 1B), consistent with our previous finding that PAS-B is critical for new pole localization *in vivo* (11). We also previously showed that cdG binds in PAS-B of CckA, stimulating phosphatase activity (11). While most PAS domains typically sense only one ligand (19, 32), our finding that CckA PAS-B responds to both cdG and DivL adds to a small set of PAS domains that interact with multiple structurally unrelated ligands (19,33, 34). A recent report identified a point mutant within the CckA PAS-B domain that led to increased expression of a subset of early predivisional, CtrA-dependent genes (35), suggesting that this mutant may specifically affect CckA interaction with DivL, cdG, or another signal, and that this mutant’s effect is restricted to the early predivisional cell when CckA first becomes active as a kinase.

Consistent with Childers et al. (17), our data suggest that different conformations of the DivL PAS domains can communicate with PAS-B of CckA to regulate catalytic function (Fig 1B). This manner of regulation differs from the more common paradigm of PAS-mediated signaling, in which a given signal simply promotes one conformation of the target kinase (19). Rather, DivL can actively promote both the kinase and phosphatase activities of CckA, presumably through different contacts between CckA PAS-B and some part of the DivL PAS domains. Our truncation analysis of the DivL PAS domains (Fig 2C) and prior studies indicate that its PAS domains B-D are critical domains involved in CckA regulation, but future investigations will be necessary to determine which domain(s) of DivL directly communicate with CckA and whether DivL also parses multiple input signals to determine its subcellular regulation of CckA activity.

Our data indicate that oligomerization is important for CckA kinase but not phosphatase activity, as only kinase activity is surface density dependent (11). The CckA PAS-A domain, a key driver of oligomerization, is necessary for stimulation of kinase but not phosphatase activity by DivL (Fig. 3). In most HKs, a signal in the PAS domain torques the central alpha-helical spine of the protein to regulate downstream catalytic domains (20, 36). The proline-rich linker between PAS-A and PAS-B of CckA is inconsistent with this alpha-helix, but it may provide a semi-rigid connection still capable of torque. Alternatively, oligomerization via PAS-A and RD (Fig. 4) may regulate kinase activity on the conserved principle of stabilizing a rotation of the ATP binding domain relative to the active site histidine by changing CckA-CckA contacts between higher and lower assembly states. Indeed, structural evidence shows that binding of ATP versus ADP in the CA domain is sufficient to switch an HK between kinase and phosphatase conformations, respectively (25). Broadly, this demonstrates that the molecular impetus for switching between catalytic functions may come from many different parts of the protein, and oligomerization could provide multiple contact points to regulate kinase activity (Fig 1B).

Further, oligomerization offers a straightforward mechanism for linking kinase activity to accumulation of a protein within a subcellular niche. In *Caulobacter*, a specialized collection of signaling and structural proteins at the new cell pole coordinates the stimulation of CckA kinase activity with polar organelle biogenesis (Fig. 5). The polar matrix protein PopZ defines the boundaries of DivL and CckA polar accumulation, and consequently the surface density of CckA, at the new pole (22, 37–39). Thus, the boundaries of the PopZ microdomain give subcellular context to CckA’s density-dependent kinase activity.

Additional layers of regulation may promote CckA kinase activity at the new cell pole. Tsokos et al. proposed that the new pole constitutes a ‘protected zone’ in which other factors cannot inhibit CckA kinase activity (16). Moreover, the DivL mutant A601L directly stimulates CckA kinase activity, and suggests that additional factors at the new cell pole, reviewed elsewhere (40), may be necessary to promote kinase activity of the DivL-CckA complex *in vivo.* Our study of the DivL-CckA interaction reveals that DivL can switch between directly stimulating CckA kinase and phosphatase activities, that CckA parses multiple signals through its PAS-B domain, and that CckA oligomerization is critical for its kinase activity. Thus, a multi-sensor domain architecture provides a means for integrating a complex array of input signals within the different spatial and temporal contexts required for a single master regulator that determines divergent cell fates.

## Materials and Methods

### Cloning, protein expression, and purification

PAS domain cut-offs in CckA and DivL were assigned using the HHpred protein homology web server (23, 24) and described in detail previously (11). New plasmids for this study were created using Gibson assembly as designed through the J5 cloning system(45). Other plasmids were described previously (11, 17, 28, 46, 47). Plasmids were transformed into *E. coli* via heat shock. A complete list of plasmids and strains used in this study is available in Supplementary Table 2. CckA and DivL variants were expressed and purified as described previously (11, 17, 47), and are described at length in the Supplementary Methods.

### Gel filtration chromatography

Protein purification beyond Ni-NTA affinity was performed via gel filtration chromatography using a GP-250 gradient programmable chromatographer. Concentrated samples were separated using a Superdex 200 10/300 GL column. Fractions of 0.3 mL were collected, analyzed, and pooled to optimize purity.

Studies of CckA oligomerization via analytical gel filtration chromatography were performed using a GE Healthcare Superdex 200 Increase 10/300 GL column and a Bio-Rad NGC chromatography system. Apparent molecular weights of CckA oligomeric complexes were assigned using a standard curve based on elution volumes of a Bio-Rad premixed gel filtration standard (cat #151-1901). Elution was performed at 0.35 mL/min. CckA samples of approximately 150 μL were loaded at approximately 100 and eluted at 350 μL/min. A complete list of SEC experiments is given in Supplementary Table S1.

### Production of Large Unilamellar Liposomes

Liposomes were made as described previously (11), with some adjustments. A mixture of 900μL of 10 mg/mL 1,2-dioleoyl-*sn*-glycero-3-phospho-(1′-rac-glycerol) (sodium salt) (DOPG: product 840475), was mixed with 1 mL of 1 mg/mL 1,2-dioleoyl-*sn*-glycero-3-[(N-(5-amino-1-carboxypentyl)iminodiacetic acid)succinyl] (nickel salt) (DGS-NTA(Ni): product 790404C). Both lipids contain two 18-carbon acyl chains with a c/s-alkene between carbon atoms 9 and 10 in each chain to membrane fluidity at room temperature. The chloroform solutions were well mixed in a glass scintillation vial, and the solvent was evaporated under a gentle stream of nitrogen for 1 hour to leave a clear film with no clumps. The film was then rehydrated in 500 μL to 20 mg/mL in liposome rehydration buffer (100 mM KCl, 20 mM HEPES-KOH pH 8.0). The buffer was vigorously resuspended with a pipette until all of the lipid was dissolved. The vial containing the lipid mixture was then subjected to 10 freeze/thaw cycles in liquid nitrogen and a 37°C water bath. The freeze-thaw mixture was then extruded for 11 passes through 100 nm pores of a polycarbonate filter using the Avanti Mini-Extruder. Following extrusion, the liposomes were diluted in equal volume ultrapure water (to a final concentration of 10 mg/mL) and aliquots were stored under nitrogen in plastic tubes to extend shelf life.Production of large unilamellar vesicles has also been described previously (48–50).

### Radiolabeling phosphorylation assays

Radiolabeled auto-phosphorylation assays were similar to previous protocols (11). CckA constructs (5 μM) were incubated in low salt kinase buffer (50 mM KCl, 10 mM HEPES-KOH pH 8.0) with 5 mM MgCl_2_ and 0.5 mM ATP in 25 μL reaction volumes. Glycerol was removed from protein samples via overnight dialysis prior to reactions to match the solvent conditions within the liposome lumens. Owing to opposite ionic strength preferences for kinase activity as compared to WT CckA, kinase assays for CckA ΔPAS-B and ΔPAS-AB were conducted in high salt buffer (200 mM KCl and 50 mM HEPES-KOH pH 8.0). For Figure 4B, CckA ΔPAS-B conditions are normalized to the no-DivL comparison in each case. For all LUV-based assays, CckA and DivL were allowed to incubate for 10 minutes with varying amounts of LUVs prior to addition of ATP stocks. Kinase assays on liposomes were conducted at a surface density of 350 molecules per liposome of CckA, with 350 molecules per liposome of DivL where appropriate, unless otherwise noted. Maximum density corresponds to 1100 molecules per liposome.

Radiolabeled ATP was supplemented at of 2 μCi [γ-^32^P] ATP per reaction. Reactions were quenched after 3 minutes in 2x Laemmli sample buffer, and the quenched reaction mixtures were loaded onto 4-15% gradient polyacrylamide gels and subjected to electrophoresis. Alternatively, at this step for titration curve experiments, we used a nitrocellulose dot blot assay to separate phosphorylated protein from the reaction mixture, described previously (11). The extent of auto-phosphorylation was measured by exposing a phosphor screen to the gels for at least 3 hours, and the screen was subsequently imaged on aTyphoon storage phosphorimager (Molecular Dynamics). Band intensities were quantified using ImageJ.

For phosphatase assays, CckA was allowed to auto-phosphorylate in solution, and CckA~P was subsequently purified away from ATP. Depletion of ATP was accomplished by first rapidly passing the crude reaction mix through a desalting column to remove most of the nucleotides. Hexokinase (12.5 units) and glucose (10 mM) were then added to convert any remaining ATP to ADP over 10 minutes. CckA~P was then deposited on liposomes that did or did not contain equimolar DivL to test whether DivL impacts CckA dephosphorylation. For phosphatase assays supplemented with ADP, the column purification step was skipped, permitting complete conversion of ATP to ADP by hexokinase.

## Acknowledgements

We thank Seth Childers, Keith Moffat, Kim Kowallis, Sam Duvall, Michael Collins, and Keren Lasker for helpful discussion and critical review of the manuscript, and to Saumya Saurabh for helpful improvements in the liposome preparation protocol. Research reported in this publication was supported by the National Institute of General Medical Sciences of the National Institute of Health under award numbers: T32 GM007276, supporting T.H.M., as well as NIH award numbers R01 GM032506 and R35-GM118071 to L.S. The content is solely the responsibility of the authors and does not necessarily represent the official view of the National Institute of Health. L.S. is a Chan Zuckerberg Biohub Investigator.

## References

1. Shapiro L, McAdams HH, Losick R (2009) Why and how bacteria localize proteins. Science 326(5957):1225–8.

2. Quon KC, Marczynski GT, Shapiro L (1996) Cell cycle control by an essential bacterial two-component signal transduction protein. Cell 84(1):83–93.

3. Laub MT, Chen SL, Shapiro L, McAdams HH (2002) Genes directly controlled by CtrA, a master regulator of the Caulobacter cell cycle. Proc Natl Acad Sci U S A 99(7):4632–7.

4. Quon KC, Yang B, Domian IJ, Shapiro L, Marczynski GT (1998) Negative control of bacterial DNA replication by a cell cycle regulatory protein that binds at the chromosome origin. Proc Natl Acad Sci U S A 95:13600–13605.

5. Domian IJ, Quon KC, Shapiro L (1997) Cell type-specific phosphorylation and proteolysis of a transcriptional regulator controls the G1-to-S transition in a bacterial cell cycle. Cell 90(3):415–24.

6. Jacobs C, Domian IJ, Maddock JR, Shapiro L (1999) Cell cycle-dependent polar localization of an essential bacterial histidine kinase that controls DNA replication and cell division. Cell 97(1):111–20.

7. Jacobs C, Ausmees N, Cordwell SJ, Shapiro L, Laub MT (2003) Functions of the CckA histidine kinase in Caulobacter cell cycle control. Mol Microbiol 47(5):1279–1290.

8. Heinrich K, Sobetzko P, Jonas K (2016) A Kinase-Phosphatase Switch Transduces Environmental Information into a Bacterial Cell Cycle Circuit. PLOS Genet 12(12):e1006522.

9. Vo CD, et al. (2017) Repurposing Hsp90 inhibitors as antibiotics targeting histidine kinases. Bioorg Med Chem Lett 27:5235–5244.

10. Iniesta AA, Hillson NJ, Shapiro L (2010) Cell pole-specific activation of a critical bacterial cell cycle kinase. Proc Natl Acad Sci 107(15):7012–7017.

11. Mann TH, Childers WS, Blair JA, Eckart MR, Shapiro L (2016) A cell cycle kinase with tandem sensory PAS domains integrates cell fate cues. Nat Commun 7:1–12.

12. Biondi EG, et al. (2006) Regulation of the bacterial cell cycle by an integrated genetic circuit. Nature 444(7121):899–904.

13. Lori C, et al. (2015) Cyclic di-GMP acts as a cell cycle oscillator to drive chromosome replication. Nature 523:236–239.

14. Christen M, et al. (2010) Asymmetrical distribution of the second messenger c-di-GMP upon bacterial cell division. Science328(5983):1295–7.

15. Wu J, Ohta N, Zhao J-L, Newton A (1999) A novel bacterial tyrosine kinase essential for cell division and differentiation. Proc Natl Acad Sci 96(23):13068–13073.

16. Tsokos CG, Perchuk BS, Laub MT (2011) A dynamic complex of signaling proteins uses polar localization to regulate cell-fate asymmetry in Caulobacter crescentus. Dev Cell 20(3):329–41.

17. Childers WS, et al. (2014) Cell fate regulation governed by a repurposed bacterial histidine kinase. PLoS Biol 12(10):e1001979.

18. Reisinger SJ, Huntwork S, Viollier PH, Ryan KR (2007) DivL performs critical cell cycle functions in Caulobacter crescentus independent of kinase activity. J Bacteriol 189(22):8308–20.

19. Möglich A, Ayers RA, Moffat K (2009) Structure and signaling mechanism of Per-ARNT-Sim domains. Structure 17(10):1282–94.

20. Diensthuber RP, Bommer M, Gleichmann T, Möglich A (2013) Full-length structure of a sensor histidine kinase pinpoints coaxial coiled coils as signal transducers and modulators. Structure 21(7):1127–36.

21. Westbye AB, et al. (2018) The protease ClpXP and the PAS-domain protein DivL regulate CtrA and gene transfer agent production in Rhodobacter capsulatus. Appl Environ Microbiol (April):AEM.00275–18.

22. Lasker K, et al. (2017) Phospho-signal flow from a pole-localized microdomain spatially patterns transcription factor activity. bioRxiv.

23. Söding J, Biegert A, Lupas AN (2005) The HHpred interactive server for protein homology detection and structure prediction. Nucleic Acids Res 33(SUPPL. 2):244–248.

24. Zimmermann L, et al. (2017) A Completely Reimplemented MPI Bioinformatics Toolkit with a New HHpred Server at its Core. J Mol Biol (Table 1):1–7.

25. Casino P, Miguel-Romero L, Marina A (2014) Visualizing autophosphorylation in histidine kinases. Nat Commun 5:3258.

26. Huynh TN, Stewart V (2011) Negative control in two-component signal transduction by transmitter phosphatase activity. Mol Microbiol 82(2):275–286.

27. Dubey BN, et al. (2016) Cyclic di-GMP mediates a histidine kinase / phosphatase switch by noncovalent domain cross-linking. Sci Adv 2(September):1–10.

28. Chen YE, Tsokos CG, Biondi EG, Perchuk BS, Laub MT (2009) Dynamics of two Phosphorelays controlling cell cycle progression in Caulobacter crescentus. J Bacteriol 191(24):7417–29.

29. Little R, Martinez-Argudo I, Perry S, Dixon R (2007) Role of the H domain of the histidine kinase-like protein NifL in signal transmission. J Biol Chem 282(18):13429–13437.

30. Eswaramoorthy P, Guo T, Fujita M (2009) In vivo domain-based functional analysis of the major sporulation sensor kinase, KinA, in Bacillus subtilis. J Bacteriol 191(17):5358–5368.

31. Willett JW, Crosson S (2017) Atypical modes of bacterial histidine kinase signaling. Mol Microbiol 103(2):197–202.

32. Henry JT, Crosson S (2011) Ligand-Binding PAS Domains in a Genomic, Cellular, and Structural Context. Annu Rev Microbiol 65(1):261–286.

33. Scheuermann TH, et al. (2009) Artificial ligand binding within the HIF2alpha PAS-B domain of the HIF2 transcription factor. Proc Natl Acad Sci U S A 106(2):450–5.

34. Hoffman EC, et al. (1991) Cloning of a factor required for activity of the Ah (dioxin) receptor. Science 252(5008):954–958.

35. Narayanan S, Kumar L, Radhakrishnan SK (2018) Sensory domain of the cell cycle kinase CckA in Caulobacter crescentus regulates the differential DNA binding activity of the master regulator CtrA. bioRxiv.

36. Möglich A, Ayers R a., Moffat K (2010) Addition at the Molecular Level: Signal Integration in Designed Per-ARNT-Sim Receptor Proteins. J Mol Biol 400(3):477–486.

37. Ebersbach G, Briegel A, Jensen GJ, Jacobs-Wagner C (2008) A self-associating protein critical for chromosome attachment, division, and polar organization in caulobacter. Cell134(6):956–68.

38. Bowman GR, et al. (2008) A Polymeric Protein Anchors the Chromosomal Origin/ParB Complex at a Bacterial Cell Pole. Cell 134(6):945–955.

39. Holmes JA, et al. (2016) Caulobacter PopZ forms an intrinsically disordered hub in organizing bacterial cell poles. Proc Natl Acad Sci 113(44):12490–12495.

40. Lasker K, Mann TH, Shapiro L (2016) An intracellular compass spatially coordinates cell cycle modules in Caulobacter crescentus. Curr Opin Microbiol 33:131–139.

41. Chen YE, Tropini C, Jonas K, Tsokos CG, Huang KC (2010) Spatial gradient of protein phosphorylation underlies replicative asymmetry in a bacterium. Proc Natl Acad Sci U S A.

42. Viollier PH, Sternheim N, Shapiro L (2002) Identification of a localization factor for the polar positioning of bacterial structural and regulatory proteins. Proc Natl Acad Sci U S A 99(21):13831–6.

43. Matroule J-Y, Lam H, Burnette DT, Jacobs-Wagner C (2004) Cytokinesis monitoring during development; rapid pole-to-pole shuttling of a signaling protein by localized kinase and phosphatase in Caulobacter. Cell 118(5):579–90.

44. Sanselicio S, Bergé M, Théraulaz L, Radhakrishnan SK, Viollier PH (2015) Topological control of the Caulobacter cell cycle circuitry by a polarized single-domain PAS protein. Nat Commun 6(May):7005.

45. Hillson NJ, Rosengarten RD, Keasling JD (2012) j5 DNA Assembly Design Automation. ACS Synth Biol 1116:14–21.

46. Rocco CJ, Dennison KL, Klenchin V a., Rayment I, Escalante-Semerena JC (2008) Construction and use of new cloning vectors for the rapid isolation of recombinant proteins from Escherichia coli. Plasmid 59(3):231–237.

47. Blair JA, et al. (2013) Branched signal wiring of an essential bacterial cell-cycle phosphotransfer protein. Structure 21(9):1590–601.

48. Hope MJ, Bally MB, Webb G, Cullis PR (1985) Production of large unilamellar vesicles by a rapid extrusion procedure: characterization of size distribution, trapped volume and ability to maintain a membrane potential. Biochim Biophys Acta 812:55–65.

49. Mayer LD, Hope MJ, Cullis PR (1986) Vesicles of variable sizes produced by a rapid extrusion procedure. Biochim Biophys Acta 858:161–168.

50. Kim J, Blackshear PJ, Johnson JD, McLaughlin S (1994) Phosphorylation reverses the membrane association of peptides that correspond to the basic domains of MARCKS and neuromodulin. Biophys J 67(1):227–37.

